# A near telomere-to-telomere phased reference assembly for the male mountain gorilla

**DOI:** 10.1101/2024.10.28.620258

**Authors:** David R. Nelson, Richard Muvunyi, Khaled M. Hazzouri, Jean-Claude Tumushime, Gaspard Nzayisenga, Nziza Julius, Wim Meert, Latifa Karim, Wouter Coppieters, Katherine M. Munson, DongAhn Yoo, Evan E. Eichler, Kourosh Salehi-Ashtiani, Jean-Claude Twizere

## Abstract

The endangered mountain gorilla, Gorilla beringei beringei, faces numerous threats to its survival, highlighting the urgent need for genomic resources to aid conservation efforts. Here, we present a near telomere-to-telomere, haplotype-phased reference genome assembly for a male mountain gorilla generated using PacBio HiFi (26.77× ave. coverage) and Oxford Nanopore Technologies (52.87× ave. coverage) data. The resulting non-scaffolded assembly exhibits exceptional contiguity, with contig N50 of ∼95 Mbp for the combined pseudohaplotype (3,540,458,497 bp), 56.5 Mbp (3.1 Gbp) and 51.0 Mbp (3.2 Gbp) for each haplotype, an average QV of 65.15 (error rate = 3.1 × 10-7), and a BUSCO score of 98.4%. These represent substantial improvements over most other available primate genomes. This first high-quality reference genome of the mountain gorilla provides an invaluable resource for future studies on gorilla evolution, adaptation, and conservation, ultimately contributing to the long-term survival of this iconic species.

## Background & Summary

The mountain gorilla (*Gorilla beringei beringei*, see Figure 1) is an endangered subspecies of the eastern gorilla, with a population of approximately 1,063 individuals remaining in the wild as of 2018^1^. These great apes are found exclusively in the high-altitude forests of the Virunga Massif, Sarambwe Reserve and Bwindi Impenetrable National Parks, spanning the borders of Rwanda, Uganda, and the Democratic Republic of Congo, at elevations ranging from 1,100 to 4,500 meters above sea level^2^. As one of our closest living relatives, sharing approximately 98% of their DNA with humans^2^, the mountain gorilla holds deep evolutionary, ecological, and conservation importance. Understanding their genome is crucial for deciphering the genetic basis of their unique adaptations, such as their ability to thrive at high altitudes, their population history, and susceptibility to diseases, as well as for informing conservation strategies to protect this endangered species. Obtaining high-quality genomic samples from mountain gorillas is exceptionally challenging due to their limited population size, remote habitats, and strict conservation regulations^3^. Furthermore, until recently, producing enough long-read sequencing data for high-quality mammalian genomes has been costly, limiting candidate species for genomic projects. These factors have hindered the generation of a comprehensive reference genome for this subspecies. Previous genomic studies on mountain gorillas generally used sequencing reads from *G. beringei beringei* to align with other reference genomes, including the Eastern lowland gorilla (*Gorilla beringei graueri*)^4^, limiting the scope of genetic analyses and comparative studies. A near telomere-to-telomere (T2T) phased reference assembly would provide a cutting-edge resource for investigating the complex evolutionary history, population dynamics, and genetic diversity of mountain gorillas, similar to recent advances in human genomics^5^.

**Figure 1:**
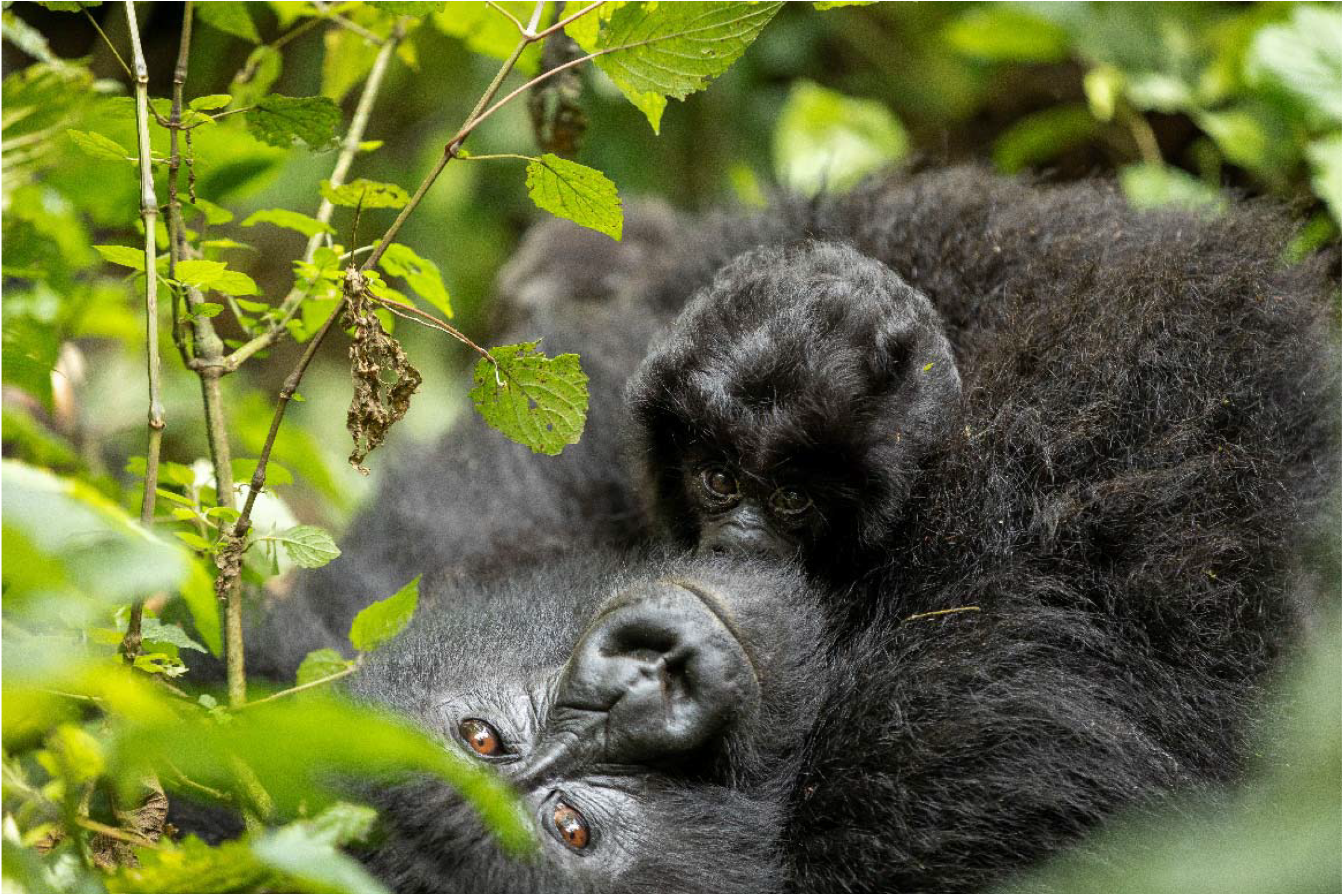
Male mountain gorilla (*Gorilla beringei beringei*) named Igicumbi, donor of blood sample used for DNA extraction for sequencing, photographed with the mother. Opportunities for biological specimen sample collection in these endangered animals is typically restricted to periods where human intervention may be necessary to prevent life-threatening conditions. The two-year-old gorilla, Igicumbi (13.4 kg), underwent critical veterinary intervention, allowing for an opportunistic blood sampling for samples with high molecular weight (HMW) DNA extraction.

Recent advancements in long-read sequencing technologies, such as Pacific Biosciences (PacBio; Menlo Park, CA, USA) HiFi (high-fidelity) and Oxford Nanopore Technologies (ONT; Oxford, UK), have revolutionized the field of genomics by enabling the generation of highly contiguous and accurate genome assemblies^6^. These technologies, combined with state-of-the-art assembly algorithms and phasing methods, have made it possible to construct near-T2T assemblies for various species, including humans^7^ and other great apes^8,9^. However, the application of these technologies to the mountain gorilla genome has been limited by the scarcity of high-quality samples and the challenges associated with obtaining them from wild populations^10^.

We present the first near-T2T phased reference assembly for the male mountain gorilla, overcoming the challenges associated with obtaining high-quality genomic samples from this endangered subspecies. We used a combination of long-read sequencing technologies and advanced assembly and k-mer-based phasing approaches to sequence the *G. beringei beringei* genome. By leveraging the power of PacBio HiFi reads, which offer both long read lengths (15-20 kbp) and high accuracy (>99.9%)^11^, and ONT ultra-long reads, which can span hundreds of kilobases^12^, we were able to generate accurate haplotype separation in the absence of parental information.

We constructed highly contiguous and accurate assemblies that capture the majority of the mountain gorilla genome, including complex regions such as centromeres and telomeres (Table 1, Figure 2). Compared to the T2T *G. gorilla* assembly^9^, we find that ∼90% of each of the chromosomes align with as few as two contigs (Figure 4 and Table 2). The resulting assembly exhibits exceptional contiguity, with contig N50 of ∼95 Mbp for the combined pseudohaplotype (3.5 Gbp), 56.5 Mbp (3.1 Gbp) and 51.0 Mbp (3.2 Gbp) for each haplotype, and an average QV of 65.15 (error rate = 3.1 × 10^−7^).

**Table 1.** Comparison of assembly metrics from the new *G. beringei beringei* assemblies and three other assemblies used for comparisons. The main *G. beringei beringei assembly* has had contiguous regions integrated from each haplotype to form a singular, pseudohaplotype representative assembly^15^ for the species. Non-BUSCO^16^ metrics were generated with QUAST^17^.

| Assembly | Total length (Gbp) | GC (%) | N50 (Mbp) | L50 | BUSCOs % complete | Reference | Sequencing technology |
| --- | --- | --- | --- | --- | --- | --- | --- |
| *Gorilla gorilla* (GCF_029281585.2)<sup>13</sup> | 3.5 | 40.49 | 150 | 10 | 98.4 | NHGRI, NIH | PacBio Sequel; ONT PromethION |
| *G. beringei beringei* Joined | 3.5 | 40.59 | 95 | 14 | 98.4 | This study | PacBio Sequel; ONT PromethION |
| *G. beringei beringei* Hap 2 | 3.2 | 40.49 | 51 | 16 | 89.1 | This study | PacBio Sequel; ONT PromethION |
| *G. beringei beringei* Hap 1 | 3.1 | 40.29 | 57 | 18 | 83.1 | This study | PacBio Sequel; ONT PromethION |
| *Gorilla gorilla* (GCA_030174185.1)<sup>14</sup> (Kamilah) | 3.6 | 40.53 | 38 | 27 | 98 | UWashington | PacBio Sequel |
| *G. beringei* (GCA_963575185.1) | 2.7 | 40.87 | 0.055 | 14910 | 68.9 | IBE | Illumina |

**Figure 2:**
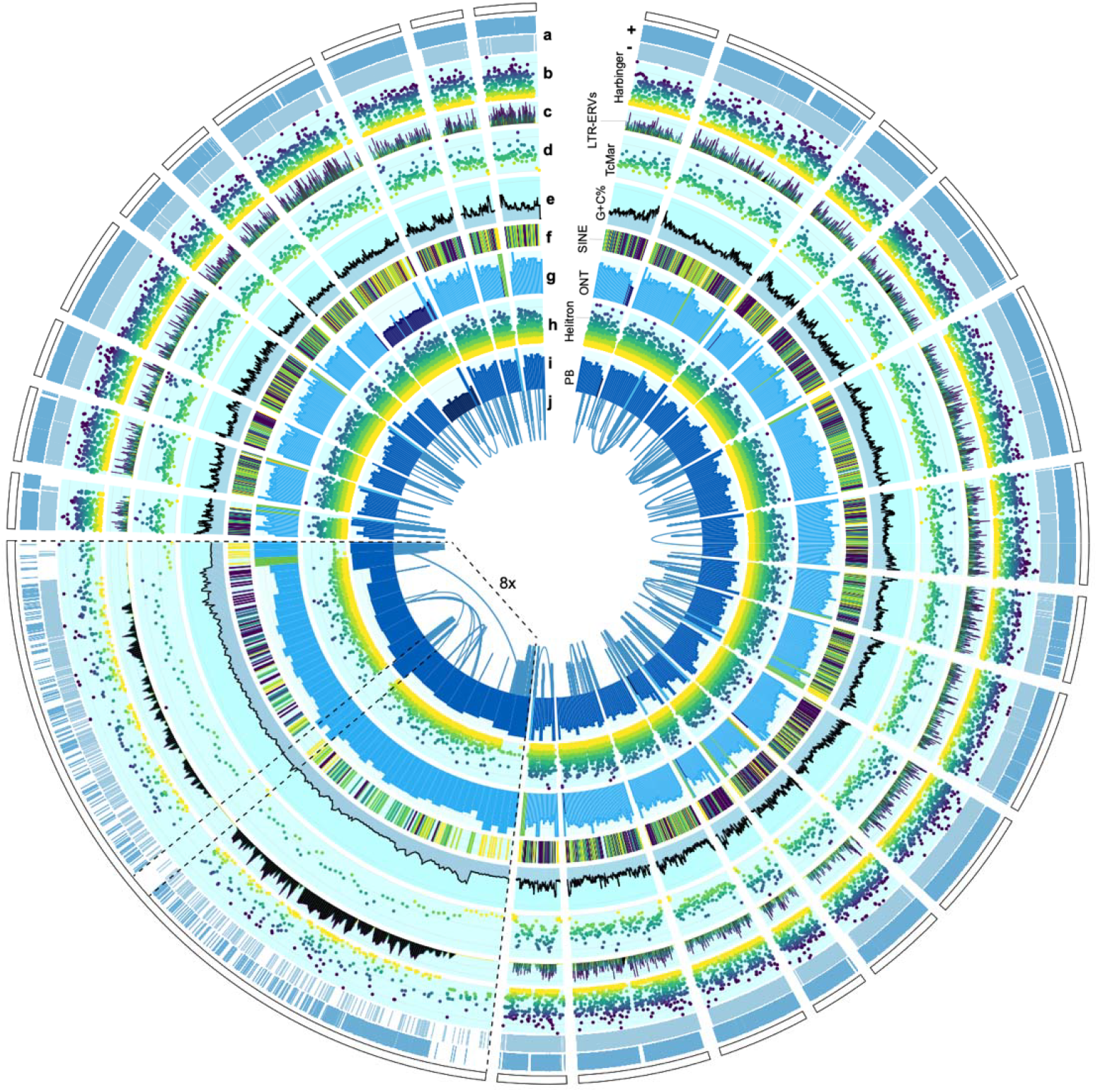
Genome map showing sequencing read coverage and annotations for the new *de novo*, near-T2T male *G. beringei beringei* genome (gberber-nt2t). The 25 largest contigs from the pseudohaplotype assembly are shown. An example T2T-sequenced chromosome is displayed at 8× zoom with the telomeres and centromere regions, identifiable by having high coverage and low gene/LTR/SINE content, bounded in dashed lines. Circos tracks represent (a) plus (top) and minus (bottom) strand *ab initio* gene predictions. (b) Harbinger elements. (c) LTR/ERV repeats, where the y-axis = % divergence from the reference. (d) TcMar elements. Repeat element tracks have a y-axis ranging from zero to 50% divergence. (e) G + C % with a y-axis ranging from 20–80 G + C %. (f) SINE elements. (g) ONT read coverage (ave. = 52.87×; see also Figure 3). (h) Helitron elements. (i) PacBio HiFi long-read coverage (ave. = 26.77×). (j) Extended tracks of high homology/similarity (i.e., identical long repeat sequences). Circos data tracks and configurations are in Dataset S4.

**Figure 3:**
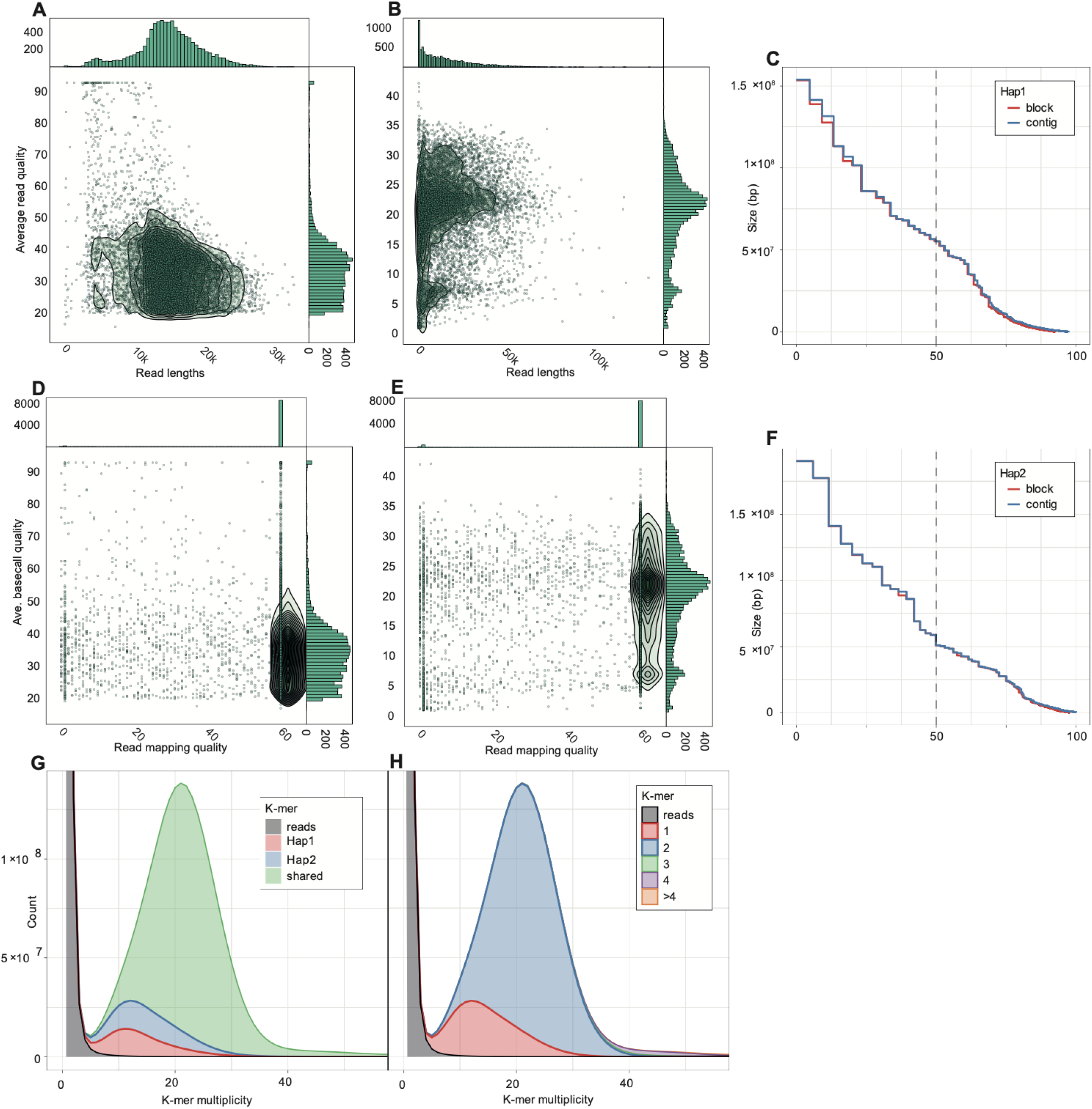
Quality assessment metrics for the phased mountain gorilla genome assembly. These results demonstrate the assembly quality and haplotype resolution across both haplotypes (Hap1 and Hap2). See Figure 4 and Table 2 for mapping of this assembly to a recent T2T *G. gorilla* genome^9^. (A,B): Sequencing read lengths (x-axis) compared to the average read quality (y-axis) for (A) PacBio HiFi and (B) ONT reads. (C,F): NG graphs: Bottom panels present NG (length-weighted median) curves for both haplotypes. The x-axis represents NG percentages, while the y-axis shows contig/block sizes. Separate curves for contigs (blue) and phased blocks (orange) are plotted. The high NG50 values and gradual curve slopes suggest excellent contiguity and completeness for both haplotypes. (D,E): Read mapping quality (QV, x-axis) compared to the average base-call quality for (D) PacBio HiFi and (E) ONT reads. QV, calculated as -10 * log10 (error rate), quantifies base-level accuracy. (G,H): K-mer spectra: These plots display k-mer frequency distributions for the Hap1 and Hap2 haplotype assemblies. The x-axis represents k-mer multiplicity (occurrence frequency), while the y-axis shows the count of distinct k-mers at each multiplicity. The bimodal distribution with peaks at 1× and 2× coverage indicates effective separation of haplotypes and high completeness.

**Figure 4:**
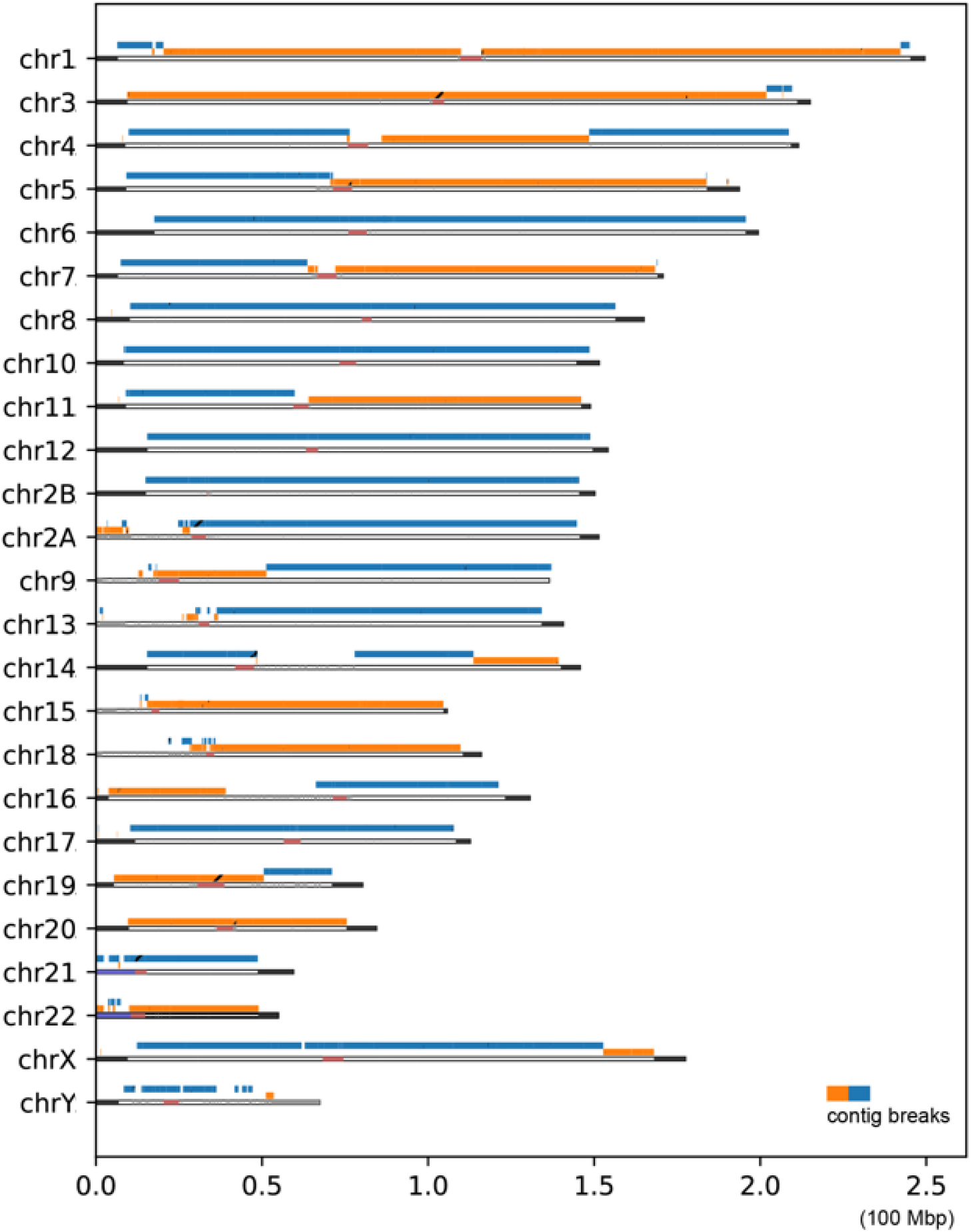
Mapping of the *G. beringei beringei* contigs to a recently published T2T *G. gorilla* genome^9^. The *G. beringei beringei* assembly aligns with an average of 2 contigs mapping per chromosome, with incomplete coverage of some satellite repeat regions (Table 2). The change of contigs is indicated by a transition of orange and blue. The T2T genomes at the bottom of each row are annotated by the following color scheme: red: centromere, purple: acrocentric p-arm, dark gray: pCht satellites, light gray: other satellites. Fig. S1 shows a zoomed-in view of Chr 1 and Table S1 shows detailed mapping statistics.

The generation of a near-T2T phased reference assembly for the mountain gorilla will be important for conservation biology, evolutionary genomics, and comparative studies among great apes. This resource will enable investigation of the genetic basis of adaptive traits, uncover the evolutionary history of mountain gorillas, and identify genomic regions associated with disease susceptibility and resilience. Moreover, the assembly will serve as a valuable reference for future population genomic studies, facilitating the development of targeted conservation strategies to protect this endangered subspecies.

## Methods

### Sample collection

Sample collection was conducted under the strict guidelines of a research permit issued by the Rwanda Development Board (RDB) for the project titled “*Obtaining high-quality genomic information and understanding the genetic diversity of mountain gorillas in Rwanda*,” obtained from the Wildlife Authority of Rwanda. A two-year-old infant male gorilla named Igicumbi exhibited lethargy and reduced feeding following his mother’s death from a severe respiratory infection. Due to his concerning condition, veterinarians conducted an immediate veterinary health intervention. After taking the animal’s body weight measurement (13.4 kg), the veterinary team performed a chemical immobilization (anesthesia). A combination of a half-dose of ketamine (23.45 mg) and dexmedetomidine (0.2 mg) was administered intramuscularly into the right thigh. The anesthesia took effect within three minutes, with full sedation achieved within five minutes. Igicumbi remained stable throughout the procedure. A physical examination was performed, and samples, including swabs and blood, were taken and stored in various sample media. The blood sample (approximately 20 mL) was drawn from the left femoral vein using BD butterfly catheter and kept in four 5 ml EDTA tubes. The samples were immediately placed in a cooler with ice packs and transported to the Rwanda Development Board Laboratory, Rubirizi, Rwanda for further analysis.

### DNA isolation and sequencing

For genomic DNA isolation, white blood cells were isolated from 5 ml of blood samples 30 hours post-collection, using a red blood cell (RBC) lysis solution (Qiagen, Hilden, Germany) and several cycles of centrifugation (2000 g at 4°C)/washes in phosphate-buffered saline (PBS, pH 7.4). Peripheral blood mononuclear cells (PMBCs) were also isolated from 5 ml of blood using the standard Percoll ® (GE Healthcare, Chicago, USA) centrifugation method. Briefly, plasma, PBMCs, and RBCs are separated following a gentle centrifugation at 350 g at room temperature, and PBMCs are washed several times with PBS to remove RBC contaminants. To isolate high molecular weight (HMW) DNA, two different kits were employed according to the manufacturers’ instructions: NucleoBond HMW DNA (Macherey-Nagel, Dueren, DE) or Monarch HMW DNA Extraction kit for tissue (New England Biolabs, Ipswich, MA, USA). DNA quality was evaluated using Qubit fluorometric quantitation (Thermo Fisher Scientific, Waltham, MA, USA) and a DNA Fragment Analyzer (5200 fragment Analyzer System, Agilent, Santa Clara, CA, USA) with an Agilent HS Genomics Kit (Agilent, DNF-468 HS Genomic DNA 50 kb Kit).

From blood samples, we generated 4,892,167 HiFi reads; of which, 3,860,717 were ‘covered’ and 2,111,243 were ‘chained’. For HiFi library preparation and sequencing, we followed the protocol described by PacBio (https://www.pacb.com/wp-content/uploads/Procedure-checklist-Preparing-whole-genome-and-metagenome-libraries-using-SMRTbell-prep-kit-3.0.pdf) with a few modifications: (i) we used Megaruptor 3 at a slower speed that gives a longer fragment length distribution: specifically, we shear once at setting 28 to set the mode length, then immediately repeat the shear program at setting 31 to reduce the long tail of molecules; (ii) we used the DNeasy PowerClean Pro cleanup kit (Qiagen, Hilden, Germany) to achieve high-purity DNA; and (iii) after library preparation, we performed a strict size selection with the PippinHT instrument (SageScience, Beverly, MA, USA) using a high-pass cutoff of 13 kbp with a 30-minute elution time (and the longer program “0.75% 9-30kb R+T 75E”) for removal of short fragments. We used a total of 4 SMRT Cells 8M on a Sequel IIe PacBio instrument.

For ONT libraries preparation and sequencing, we followed the protocol described on ONT community (https://community.nanoporetech.com/docs/prepare/library_prep_protocols/ligation-sequencing-v14-human-cfdna-multiplex-sqk-nbd114-24/v/cfm_9208_v114_revb_15may2024). Quality control of gorilla genomic DNA was performed using a Fragment Analyzer with the Agilent HS Genomics Kit (Agilent, DNF-468 HS Genomic DNA 50 kb Kit). Library preparation was conducted with approximately 200 fmol of DNA using the Ligation Sequencing Kit V14 (LSK114, ONT) following the high duplex ligation protocol. The final libraries were sequenced on a PromethION HD flow cell. We used the native barcoding Kit 24 V14, and to increase the coverage, we used four flow cells on a PromethION 2 solo instrument. Libraries were recovered after 24 hours and reloaded onto a washed flow cell for a total of three loadings over 72 hours. The total number of duplex reads was 2,479,196 (∼89 gigabytes) and the simplex reads were 6,532,927 (∼196 gigabytes). The ONT reads passing filtering were: 9,000,542 reads, of which 4,861,406 were fully corrected and 154,854 were nearly fully corrected. Overall, 1,033,384 ultra-long reads had full chains. Read and assembly QC metrics are included in the NanoPlot^18^ and Merqury^19^ analyses in the supplementary datasets.

#### Genome assembly

We used hifiasm^15^ (version 0.19.8-r603, https://github.com/chhylp123/hifiasm), a tool designed specifically for assembling long reads, to assemble the mountain gorilla genome. This approach allowed us to leverage the high accuracy and long read lengths of HiFi and ONT sequencing data to generate highly contiguous assemblies. The command was ‘hifiasm -o assembly.asm -t 32 --ul ONT_reads.fq HiFi_reads.fq’. Hifiasm generates three main outputs: two haplotype assemblies (H1 and H2) and a pseudohaplotype assembly in graph format (.gfa) which is converted to fasta with ‘awk ‘/^S/{print “>“$2;print $3}’ $LINE > “${LINE%.gfa}”.fa .’ We tested all possible kmer configurations, where i=$(sed -n “$SLURM_ARRAY_TASK_ID”p ilist.txt) \\ hifiasm -o ./k-mer-cycle/gorilla-kmer-”$i”.asm -t 7 gorilla.fq -k “$i”, and found the default k-mer assignment to be optimal for contiguity. After initial contig assembly where hifiasm assembles all reads into contigs without separating haplotypes, the heterozygous SNPs are used to phase these contigs into two sets, creating the haplotype-resolved assemblies Hap1 and Hap2, which represent the two separate chromosome sets of the diploid genome. Then, the best contig for each region is selected, based on length, quality, and consistency with surrounding areas to produce the combined, main assembly. Hifiasm uses sequences from either haplotype to fill gaps or resolve conflicts, resulting in a single, haploid-like (pseudohaplotype) representation of the genome optimized for contiguity.

### Genome quality assessment

To evaluate the quality and completeness of our assembly, we employed several tools: QUAST for evaluating assembly contiguity, Merqury (https://github.com/marbl/merqury) for k-mer-based quality assessment, SAMtools (https://github.com/samtools) for alignment statistics, NanoPlot (https://github.com/wdecoster/NanoPlot) for long-read sequencing and assembly metrics, BUSCO (https://github.com/metashot/busco) for assessing the completeness of conserved genes, and QUAST^17^ to evaluate assembly contiguity and generate standard assembly statistics. These analyses were primarily performed on the pseudohaplotype reference assembly, providing a comprehensive assessment of its quality and completeness.

Comprehensive sequencing read, assembly, and mapping statistics were generated using NanoPlot. The ONT dataset comprised 9,304,794 reads, totaling 155,097,187,007 bases, of which 150,069,675,303 bases (96.8%) were successfully aligned to the assembly. These reads demonstrated good length and moderate accuracy, with a read length N50 of 29,503 bp and a mean read length of 16,668.5 bp (standard deviation: 16,164.8 bp). The reads exhibited an average identity of 95.4% when aligned to the assembly, with a median identity of 98.5%. Read quality was variable, with a mean read quality score of 12.5 and a median of 21.4. In terms of quality distribution, 85.4% of reads (7,948,633) surpassed Q10, 76.8% (7,144,515) exceeded Q15, 59.6% (5,548,344) achieved Q20 or higher, 22.2% (2,066,884) reached Q25, and 6.3% (587,910) attained Q30 or above. This substantial set of long ONT reads complemented the HiFi dataset, contributing to the generation of our near-T2T *G. beringei beringei* assembly. The consistently high QV scores across multiplicities, with the majority of k-mers showing QV > 60, indicate superior sequence quality and low error rates in both haplotypes.

In total, PacBio HiFi reads (*n* = 4,916,103) had a read length N50 of 16,319 bp, with 74,713,988,666 bases of which 74,625,957,927 were aligned to the genome (99.9%). The reads showed an average of 99.4% identity to the assembly. The mean read length was 15,197.8, the mean read quality was 27.5, and the standard deviation of the read length was 4,775. Nearly all reads (99.7%, 4,914,842 reads) were higher than Q20, 82.4% (4,052,820 reads) were higher than Q25, and 63.8% of reads had read quality scores higher than Q30 (3,136,324 reads). The large number of high-quality HiFi and ONT reads formed the basis of the near-T2T *G. beringei beringei* assembly.

We assessed the quality of our PacBio HiFi and ONT assembled mountain gorilla genome using Merqury^19^, a reference-free method for evaluating genome assemblies. Merqury utilizes k-mer-based methods to estimate the completeness, consensus quality, and base-level accuracy of the assembly. We analyzed two haplotype assemblies (Hap1 and Hap2) separately. The analysis provided key metrics for assessing assembly quality from long reads, including quality value (QV), error rate, number of bases, and estimated number of errors. For Hap1, we obtained a QV of 65.1013 and an error rate of 3.0894 × 10^−7^. Hap2 showed a QV of 65.1958, an error rate of 3.02288 × 10^−7^. These results indicate high-quality assemblies for both haplotypes, with QV scores above 60 suggesting a high degree of base-level accuracy. The slightly lower QV and higher error rate in Hap2 may be attributed to its larger size, potentially incorporating more complex or repetitive regions of the genome. The complete Merqury output, including detailed statistics for individual contigs, was examined to identify any specific regions of lower quality that may require further attention in the assembly process.

The completeness of the gorilla genome assemblies was with regard to complete universally conserved ortholog marks was assessed using BUSCO v5.7.0^16^ with the primates_odb10 lineage dataset (created on 2024-01-08, comprising 13,780 BUSCOs from 25 genomes). The analysis was performed in eukaryotic genome mode using miniprot^20^ as the gene predictor. Results indicated a high level of completeness in the main, merged, genome, with 98.4% of BUSCOs identified as complete (97.2% single-copy and 1.2% duplicated). Only 1.1% of BUSCOs were fragmented, and 0.5% were missing. These findings suggest that the assembled genome captures the majority of expected single-copy orthologs for primates, indicating a high-quality and nearly complete assembly.

Comparison with the T2T gorilla genome was performed by aligning each of the assemblies to the T2T assembly (mGorGor1) using minimap2 (v2.26)^21^; parameters used are as follows: “*minimap2 -c -L --eqx --cs -cx asm20 -- secondary=no --eqx -Y*”. We used all maternal chromosomes and Y chromosome as the reference. The total number of contigs that mapped by alignment length >1 Mbp were counted, and the number of reference bases covered by the whole-genome alignment were quantified for each chromosome to assess relative completeness and contiguity. The alignment visualization was performed using SafFire (https://github.com/mrvollger/SafFire) and SVbyEye (https://github.com/daewoooo/SVbyEye). In addition to the high coverage previously noted, we note, however that the acrocentric p-arms (11.9 Mbp or 53.2% covered) as well as satellite sequence corresponding to the centromeres and subterminal heterochromatic caps are not yet completely assembled (29 identified out of 42 expected) in this version of the mountain gorilla genome.

### Genomic landscape of repetitive elements

For the annotation of transposable elements, we implemented a multifaceted approach. HiTE (https://github.com/CSU-KangHu/HiTE/tree/master), a fast and accurate dynamic boundary adjustment tool, was used for full-length transposable element detection and annotation in our genome assembly. We ran RepeatMasker^22^ (https://www.repeatmasker.org/) with RMblast (https://www.repeatmasker.org/rmblast/) for comprehensive repeat element identification and masking. Our findings revealed that 34.8% (1,232,080,350 bp) of the genome is composed of repetitive elements, with total interspersed repeats accounting for 29.17% (1,032,857,806 bp) of the genome sequence. Retroelements were the most abundant class of repetitive sequences, occupying 26.58% (941,219,695 bp) of the genome. This class primarily consisted of long interspersed nuclear elements (LINEs, 14.06%), short interspersed nuclear elements (SINEs, 7.70%), and long terminal repeat (LTR) elements (4.82%). LINEs, particularly L1/CIN4 elements, dominated the retroelement landscape, potentially influencing genomic plasticity and adaptive processes^23-25^. SINEs, known to play roles in gene regulation and genome evolution^26-28^, were also prevalent. LTR elements, including a substantial proportion of retroviral elements (4.53% of the genome), may contribute to genomic diversity and potentially impact immune responses. In summary, the repeat elements outlined here may influence gene regulation, genomic plasticity, and the evolution of physiological traits necessary for survival in low-oxygen environments. For instance, the abundance of LINEs and SINEs might facilitate the evolution of genes involved in oxygen transport or utilization, similar to adaptations observed in other high-altitude species^29,30^.

### Gene prediction and transcript and protein annotation

We performed *ab initio* gene prediction on the new mountain gorilla assemblies, as well as four other gorilla genome assemblies downloaded from NCBI (including one assembly consisting of solely the Y chromosome from *G. beringei beringei*). This was done using SNAP and TransDecoder. SNAP^31^ (Semi-HMM-based Nucleic Acid Parser) is a general-purpose gene finding program based on a generalized hidden Markov model, offering flexibility, speed, and accuracy in predicting gene structures across various organisms. TransDecoder, designed to identify coding regions within transcript sequences, excels at detecting open reading frames (ORFs) with high coding potential. TransDecoder’s ability to integrate protein homology evidence and identify multiple ORFs from alternatively spliced transcripts complements SNAP’s *ab initio* approach. We used SNAP with default parameters and the ‘mam39.hmm’ hidden Markov model and the latest TransDecoder Docker instance in Singularity (docker://trinityrnaseq/transdecoder) with default parameters to generate *ab initio* coding sequence and protein predictions.

### Data visualization

A suite of visualization tools was used to represent different aspects of the *G. beringei beringei* genome assembly. Mercury (https://github.com/marbl/merqury) was used to generate k-mer spectra plots and QV distribution graphs, providing visual insights into assembly completeness and accuracy. QUAST^17^ produced contiguity plots and size distribution charts, illustrating the assembly’s structural characteristics. NanoPlot created read length histograms, quality score distributions, and alignment identity plots for both PacBio HiFi and ONT reads. HiTE (https://github.com/CSU-KangHu/HiTE/tree/master) and RepeatMasker (https://www.repeatmasker.org/) with RMblast (https://www.repeatmasker.org/rmblast/) generated visualizations of transposable element distributions and divergence landscapes across the genome.

To provide a comprehensive overview of the genome assembly, we created a Circos (circos.ca) diagram using the base Circos package (http://circos.ca/). This circular plot displays various genomic elements of the near-T2T male *G. beringei beringei* genome, focusing on the 25 largest contigs, which comprise approximately 80% of the full genome. The Circos plot integrates multiple tracks of genomic information: (A) *ab initio* gene predictions on both plus (outer) and minus (inner) strands, (B) G+C content percentage (range: 20–80%), (C) LTR/ERV repeats and (D) SINE elements, both showing percentage divergence from the reference (range: 0–50%), (E) ONT read coverage, and (F) PacBio HiFi long-read coverage, with both coverage tracks ranging from 0–50×. This comprehensive visualization allows for an intuitive understanding of the genome’s structure, gene distribution, repetitive element landscape, and sequencing coverage.

## Data records

The data associated with this project is hosted at Zenodo. The sequencing reads are available in separate repositories for the ONT and HiFi reads (ONT: https://zenodo.org/records/12633418, HiFi: https://zenodo.org/records/12634532). The assemblies and genome analysis results (BUSCO, NanoPlot, Merqury, Circos) are available at doi: 10.5281/zenodo.12631957.

## Technical validation

The quality of the *G. beringei beringei* assembly was rigorously evaluated using multiple complementary approaches to ensure high accuracy, completeness, and contiguity. We assessed the base-level accuracy and completeness using Merqury, which revealed high QV for both haplotypes: the Hap1 assembly had a QV of 65.1013 and an error rate of 3.0894 × 10^−7^. Hap2 had a QV of 65.1958 and an error rate of 3.02288 × 10^−7^. The assemblies’ contiguity were evaluated using QUAST^17^, showing a contig N50 of approximately 95 Mbp for the combined pseudohaplotype (3,540,458,497 bp), and 56.5 Mbp (3.1 Gbp) and 51.0 Mbp (3.2 Gbp) for the Hap1 and Hap2 haplotype assemblies. These values represent a substantial improvement over most available primate genomes.

Further validation was performed by aligning the sequencing reads back to the assembled genome. The PacBio HiFi reads showed an average of 99.4% identity to the assembly, with 99.88% of bases successfully mapped, while 96.8% of ONT read bases were successfully aligned with an average identity of 95.4%. The high quality of the input sequencing data, with 100% of PacBio HiFi reads surpassing Q20 and 63.8% achieving Q30 or higher, further supports the reliability of our assembly. Collectively, these comprehensive validation metrics indicate that our *G. beringei beringei* genome assembly is of high accuracy, completeness, and contiguity, providing a reliable resource for future genetic and evolutionary studies of this endangered species.

Finally, the completeness of the gorilla genome assembly regarding universally conserved, single-copy markers was assessed using BUSCO v5.7.0^16^ with the primates_odb10 lineage dataset (created on 2024-01-08, comprising 13,780 BUSCOs from 25 genomes). The analysis was performed in eukaryotic genome mode using miniprot^20^ as the gene predictor. Results indicated a high level of completeness, with 98.4% of BUSCOs identified as complete (97.2% single-copy and 1.2% duplicated). Only 1.1% of BUSCOs were fragmented, and 0.5% were missing. These findings suggest that the assembled genome captures the majority of expected single-copy orthologs for primates, indicating a high-quality and nearly complete (T2T) assembly.

## Code availability

All scripts used in this work are hosted as supplementary datasets at doi:10.5281/zenodo.12631957.

## Acknowledgements

We thank the Rwanda Development Board (RDB) for authorizing this study, and the Oxford Nanopore Technologies (ONT) for their support under ORG.one project. We thank Tonia Brown for her help in copy editing the manuscript. We also thank the Genomics platform of the GIGA Institute (University of Liege), and Genomics Core of the Center for Human Genetics (KU Leuven). This work was supported by the Académie de Recherche et d’Enseignement Supérieur – Commission de la Coopération au Développement (ARES-CCD, Belgium), ARES-PRD2024-Twizere grant and fellowships to R.M and J-C.Tumushime. This research was also supported by NYUAD Faculty Research Funds (AD060) from the Division of Science (NYUAD, UAE). J-C. Twizere is a Senior Investigator of the Fonds de la Recherche Scientifique (FRS-FNRS). This research was supported, in part, by funding from the National Institutes of Health (NIH) grant R01 HG002385 to E.E.E. E.E.E. is an investigator of the Howard Hughes Medical Institute.

## Author contributions

D.R.N.: Experimental design, data analysis, writing - review and editing.

R.M. : Sample collection, review and editing, and supervision. K.M.H.: Data analysis.

J-C.Tum.: Sample collection, review and editing.

G.N. : Sample collection.

J.N.: Sample collection, review and editing, and supervision.

W.M.: Methodology HMW DNA preparation, library preparation, and PacBio HiFi sequencing.

L.K.: HMW DNA prepartion, library preparation, and ONT sequencing.

W.C.: ONT Data analysis,and supervision.

K.M.M.: Methodology, review and editing.

D.Y.: Data analysis, review and editing.

E.E.E.: Data analysis, review and editing, supervision, funding acquisition.

K.SA.: Experimental design, review and editing, supervision, funding acquisition.

J-C.Tw.: Experimental design, review and editing, supervision, funding acquisition.

## Notes

https://zenodo.org/records/12631957

https://zenodo.org/records/12634532

https://zenodo.org/records/12633418

### Competing Interest Statement

The authors have declared no competing interest.

