## supplemental information for "A near telomere-to-telomere phased reference assembly for the male mountain gorilla"

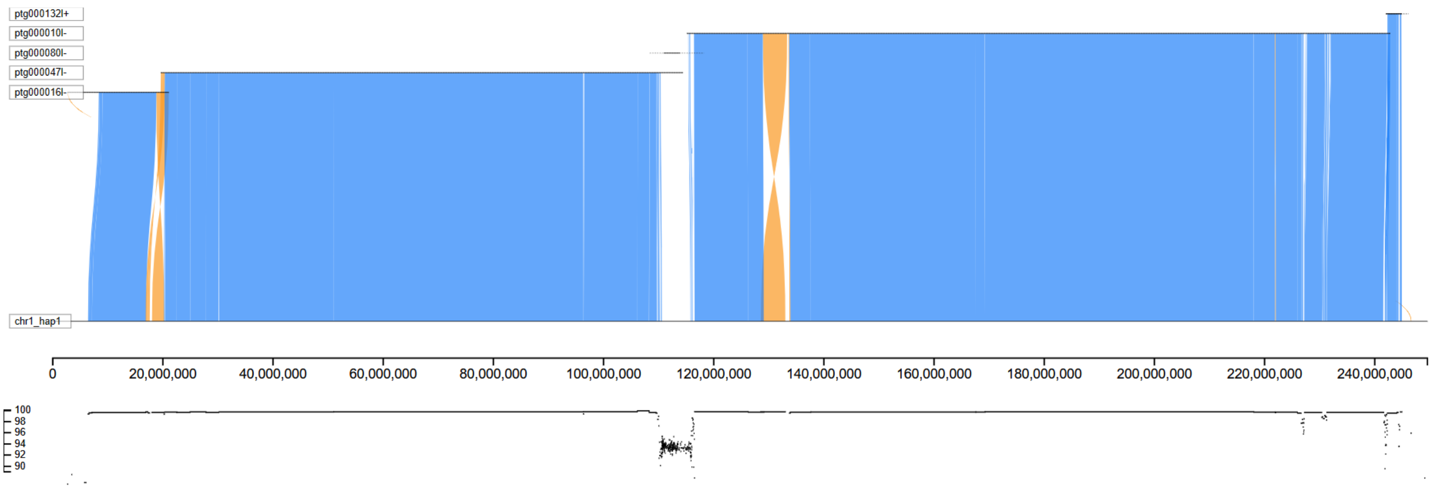


**Figure S1**: **Chromosome 1 comparison.** Sapphire view showing Minimap [1] alignments between four contigs from the *G. beringei beringei* assembly to a recently released T2T *G. gorilla* genome [2] (GCF_029281585.2). Syntenic regions in the same (blue) or inverted (orange) orientation are shown. Below the chromosomal coordinates (in bp) is shown the % identity. The centromere HOR region is delineated by a series of more divergent alignments.

| T2T- Gorilla[2] chrom (maternal + chrY) | Pseudohaplotype | | | haplotype 1 | | | haplotype 2 | | |
|---|---|---|---|---|---|---|---|---|---|
| | # of contigs | Coverage (bp) | % of reference covered | # of contigs | Coverage (bp) | % of reference covered | # of contigs | Coverage (bp) | % of reference covered |
| chr1 | 4 | 237,707,687 | 95.24% | 14 | 180,940,045 | 72.49% | 4 | 231,546,192 | 92.77% |
| chr2A | 1 | 137,378,233 | 90.72% | 10 | 112,909,482 | 74.56% | 1 | 132,178,282 | 87.29% |
| chr2B | 1 | 136,062,712 | 90.56% | 4 | 125,838,800 | 83.75% | 6 | 128,478,292 | 85.51% |
| chr3 | 2 | 200,939,185 | 93.43% | 2 | 177,378,364 | 82.47% | 5 | 195,416,211 | 90.86% |
| chr4 | 3 | 204,062,820 | 96.47% | 3 | 193,087,438 | 91.28% | 5 | 196,609,431 | 92.94% |
| chr5 | 3 | 185,082,376 | 95.53% | 6 | 161,707,587 | 83.46% | 7 | 152,792,470 | 78.86% |
| chr6 | 1 | 190,508,696 | 95.52% | 12 | 158,865,908 | 79.65% | 1 | 187,323,059 | 93.92% |
| chr7 | 4 | 170,487,976 | 99.82% | 10 | 137,133,811 | 80.29% | 5 | 165,073,671 | 96.65% |
| chr8 | 1 | 155,108,187 | 93.96% | 1 | 149,650,825 | 90.66% | 4 | 154,695,553 | 93.71% |
| chr9 | 2 | 124,320,708 | 90.56% | 2 | 122,796,978 | 89.45% | 6 | 86,568,840 | 63.06% |
| chr10 | 1 | 141,277,892 | 93.21% | 6 | 132,036,850 | 87.12% | 1 | 140,296,339 | 92.57% |
| chr11 | 2 | 138,838,189 | 93.25% | 4 | 134,675,392 | 90.46% | 5 | 115,150,124 | 77.34% |
| chr12 | 1 | 136,227,336 | 88.33% | 1 | 136,261,006 | 88.35% | 4 | 136,381,984 | 88.43% |
| chr13 | 2 | 116,586,957 | 82.81% | 5 | 101,931,986 | 72.40% | 7 | 108,227,660 | 76.87% |
| chr14 | 4 | 103,190,271 | 70.76% | 4 | 98,010,849 | 67.21% | 8 | 100,928,580 | 69.21% |
| chr15 | 1 | 89,872,724 | 84.96% | 2 | 84,764,302 | 80.13% | 10 | 72,861,251 | 68.87% |
| chr16 | 2 | 124,758,120 | 95.44% | 11 | 63,326,728 | 48.44% | 3 | 112,952,899 | 86.41% |
| chr17 | 1 | 109,562,186 | 97.19% | 10 | 91,383,663 | 81.06% | 1 | 106,719,769 | 94.67% |
| chr18 | 1 | 102,543,862 | 88.36% | 1 | 100,177,088 | 86.32% | 2 | 75,947,298 | 65.44% |
| chr19 | 3 | 61,716,027 | 76.84% | 5 | 34,808,876 | 43.34% | 6 | 46,182,474 | 57.50% |
| chr20 | 1 | 64,674,541 | 76.55% | 5 | 46,429,018 | 54.95% | 2 | 65,036,363 | 76.98% |
| chr21 | 1 | 46,498,953 | 78.14% | 2 | 30,311,096 | 50.94% | 1 | 41,473,323 | 69.69% |
| chr22 | 1 | 45,547,538 | 83.16% | 2 | 17,052,716 | 31.14% | 1 | 34,484,037 | 62.96% |
| chrX | 4 | 160,481,058 | 90.38% | 5 | 112,678,151 | 63.46% | 3 | 54,936,868 | 30.94% |
| chrY | 3 | 36,331,738 | 53.90% | 2 | 11,160,400 | 16.56% | 3 | 34,553,210 | 51.26% |
| **Average** | **2** | **128,790,639** | **87.80%** | **5.16** | **108,612,694** | **71.60%** | **4.04** | **115,072,567** | **77.79%** |

**Table S1: Alignment of *G. beringei beringei* contigs to a recently published T2T *G. gorilla* genome [2]**. The majority of primary assembly chromosomes (n=15) exhibited over 90% coverage of the T2T *G. gorilla* assembly, with an average of 2 contigs (>1 Mbp) per chromosome. This coverage falls short of 100% primarily due to long satellite repeats. As anticipated, acrocentric chromosomes (chr21, 22) and chrY, along with some satellite-rich chromosomes, demonstrated lower coverage. Alignment visualizations for each chromosome are in the supplementary data file 'SaffireViews.zip'.

1. Li, H. Minimap2: pairwise alignment for nucleotide sequences. *Bioinformatics* **2018**, *34*, 3094-3100, doi:10.1093/bioinformatics/bty191.
2. Yoo, D.; Rhie, A.; Hebbar, P.; Antonacci, F.; Logsdon, G.A.; Solar, S.J.; Antipov, D.; Pickett, B.D.; Safonova, Y.; Montinaro, F.; et al. Complete sequencing of ape genomes. *bioRxiv* **2024**, 2024.2007.2031.605654, doi:10.1101/2024.07.31.605654.
